# Molecular Crowding Tunes Material States of Ribonucleoprotein Condensates

**DOI:** 10.1101/508895

**Authors:** Taranpreet Kaur, Ibraheem Alshareedah, Wei Wang, Jason Ngo, Mahdi Muhammad Moosa, Priya R. Banerjee

**Author notes:** Corresponding author: Priya R. Banerjee.

## Abstract

Ribonucleoprotein (RNP) granules are membraneless liquid condensates that dynamically form, dissolve, and mature into a gel-like state in response to changing cellular environment. RNP condensation is largely governed by the promiscuous attractive inter-chain interactions, mediated by low-complexity domains (LCDs). Using an archetypal disordered RNP, Fused in Sarcoma (FUS), here we study how molecular crowding impacts the RNP liquid condensation. We observe that the liquid-liquid coexistence boundary of FUS is lowered by polymer crowders, consistent with the excluded volume model. With increasing bulk crowder concentration, RNP partition increases and the diffusion rate decreases in the condensed phase. Furthermore, we show that RNP condensates undergo substantial hardening wherein protein-dense droplets transition from viscous fluid to a viscoelastic gel-like state in a crowder concentration-dependent manner. Utilizing two distinct LCDs that broadly represent the most commonly occurring sequence motifs driving RNP phase transition, we reveal that the impact of crowding is largely independent of LCD charge/sequence patterns. These results are consistent with a thermodynamic model of crowder-mediated depletion interaction where inter-RNP attraction is enhanced by molecular crowding. The depletion force is likely to play key roles in tuning the physical properties of RNP condensates within a crowded intracellular space.

## Introduction

Ribonucleoprotein (RNP) granules or particles are a diverse group of subcellular compartments that are utilized by eukaryotic cells to dynamically control biomolecular processes. These nonmembrane bound assemblies, also termed as membraneless organelles (MLOs), dynamically form-dissolve and tune their physicochemical microenvironment in response to changing cellular cues[1-3]. RNP granules are enriched in proteins with low complexity domains (LCDs) that are structurally disordered[4-6], and are assumed to be formed by RNP liquid-liquid phase separation (LLPS)[7]. LLPS is a spontaneous physical process that results in the formation of co-existing liquid phases of varying densities from a homogeneous solution[2, 8]. At the molecular level, low-affinity multivalent interactions between different LCDs and their partner nucleic acids provide necessary energetic input to drive the LLPS of RNPs[9]. Experimentally, it was observed that these promiscuous interactions are tuned by several cellular physicochemical perturbations (e.g., pH, salt concentration, and non-specific interactions with biomacromolecules)[10-14].

Unlike typically utilized *in vitro* experimental conditions, *in cellulo* environments are crowded by a plethora of macromolecules that are ubiquitously present within the cellular milieu[15]. To effectively capture biomolecular dynamics in a crowded cellular environment, in vitro studies utilizing recombinant proteins employ buffer systems containing inert biocompatible polymers as crowding agents. One of the most widely used crowders is polyethylene glycol (PEG) that is a neutral hydrophilic polymer with numerous applications in crystallography, biotechnology, and medicine[16-19]. Macromolecular crowding by PEG and similar polymer agents imparts a significant excluded volume (*i.e*., space occupied by one molecule cannot be accessed by another) effect and results in alterations of molecular and mesoscale properties of biomolecules. For example, molecular crowding was shown to affect protein conformation[20-23] and RNA folding[24], conformational dynamics of intrinsically disordered proteins[25], energetics of protein self-association[26-28] and molecular recognition[29], as well as LLPS of globular proteins[30-33]. However, how macromolecular crowding alters disordered RNP condensation remains unexplored systematically.

The well-established excluded volume model of colloid-polymer mixtures predicts that the addition of a polymer chain to a neutral colloidal suspension will trigger inter-colloid attraction[34]. The underlying driving force, known as the depletion interaction, is originated due to a net entropy gain by the system via maximizing the free volume available to the polymer chains. For globular protein-crowder mixtures, this depletion interaction can induce various phase transition processes including protein crystallization[35]. In a similar vein, for RNP systems containing low complexity “sticky” domains, a considerable impact of macromolecular crowding on their phase transition is expected[36]. This idea is supported by multiple recent observations such as (i) crowding induces homotypic LLPS of nucleolar phosphoprotein Npm1 *in vitro*[37], (ii) PEG induces robust liquid phase transition of Alzheimer’s disease-linked protein Tau[38], and (iii) macromolecular crowding results in a substantial decrease in hnRNPA1 phase separation concentration[39]. However, it remains unknown whether molecular crowding impacts the fluid dynamics of RNP condensates.

The material properties of intracellular RNP granules are important determinants of their function in intracellular storage and signaling[1, 40]. Notably, a liquid-to-solid phase transition is implicated in several neurological diseases[10, 41-44]. While the roles of LCD sequence composition and charge pattern in controlling mesoscale dynamics of the RNP condensates are subjected to several recent investigations[45, 46], little is known about the effect of generalized thermodynamic forces such as crowding on RNP condensation. Here, we conduct an experimental study on the impact of macromolecular crowding on RNP liquid-liquid coexistence boundary, condensate fluidity, and transport property by RNP diffusion. Utilizing an archetypal RNP, Fused in Sarcoma (FUS), as well as representatives of the two most commonly occurring LCD sequences in eukaryotic RNPs, we demonstrate an important role of crowding in modulating the fluid dynamics of RNP condensates.

## Results

### Macromolecular crowding facilitates FUS condensation and alters droplet fluid properties

Initially, we choose to study the effects of molecular crowding on the phase behavior of FUS using PEG8000 at concentrations that mimic cellular macromolecular density (≥150 mg/ml)[47]. FUS is a stress-granule associated RNP that undergoes LLPS *in vitro* and *in cellulo* via attractive inter-protein interactions[48, 49]. Persistent FUS condensates also mature into a solid-like state during “aging” that is augmented by ALS-linked mutations[41, 50]. Therefore, FUS serves as an ideal model system for this study. Firstly, we tested the impact of crowding on the liquid-liquid coexistence boundary of the full-length FUS (FUS^FL^) in a physiologically relevant buffer (25 mM Tris, 150 mM NaCl) with varying PEG8000 concentration. We use solution turbidity in conjunction with optical microscopy to construct a phase diagram of FUS-PEG8000 mixtures, which is presented in Fig. **1a**. We observed that FUS^FL^ formed micron-scale phase-separated droplets *in vitro* at protein concentrations ≥ 2 µM without any crowding agents (Fig. **1**), consistent with the protein’s ability to undergo LLPS at physiologically relevant concentrations[50, 51]. Increasing PEG8000 concentration in the buffer solution from 0-150 mg/ml decreased FUS^FL^ phase separation concentration to < 1 µM (Fig. **1a**). These results suggest that crowding by PEG8000 facilitates FUS condensation.

**Figure-1:**
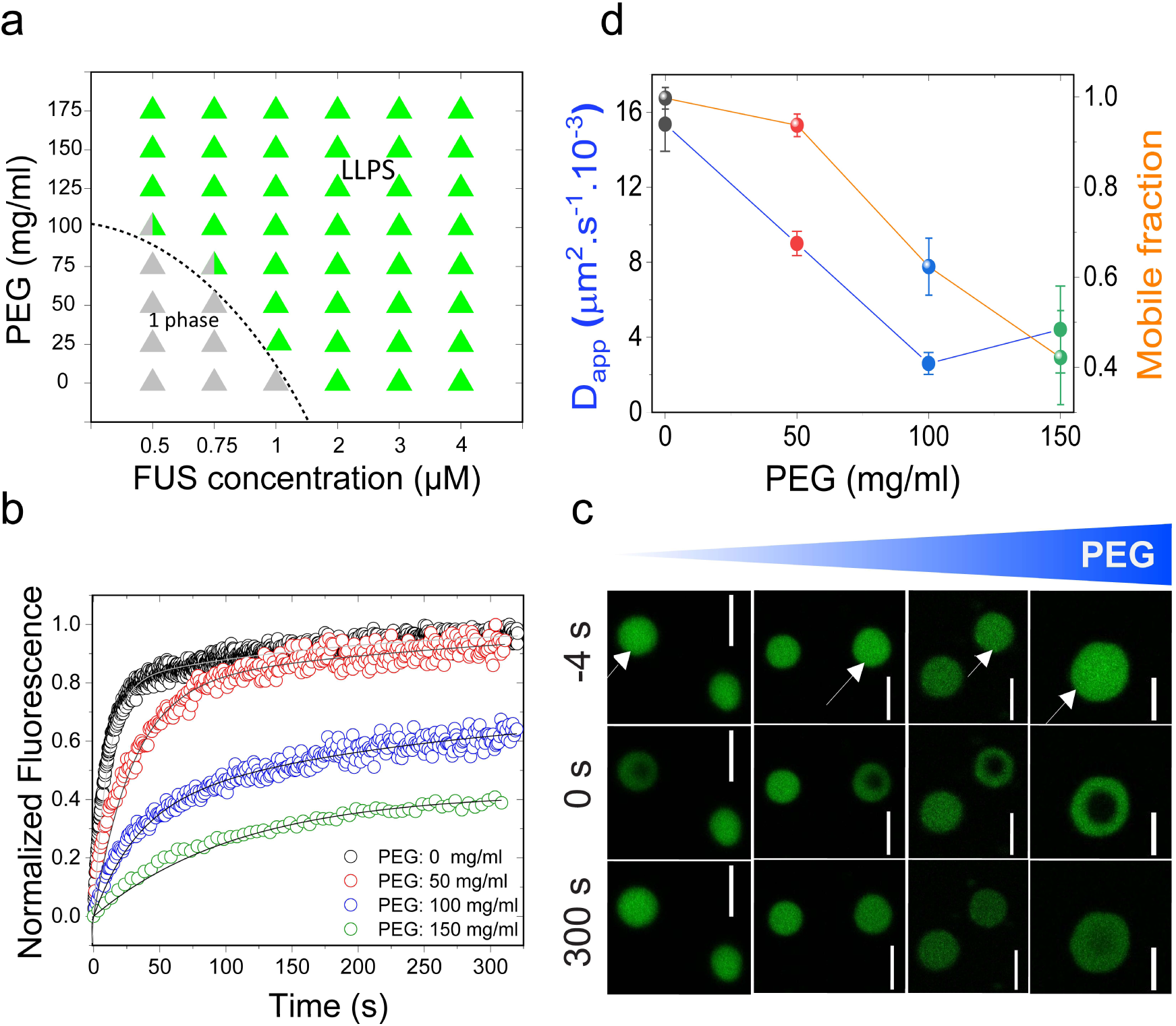
Molecular crowding facilitates FUS^FL^ liquid-liquid phase separation (LLPS) and tunes protein droplet viscoelastic properties. **(a)** Isothermal phase diagram of FUS-PEG8000 mixtures. The dotted line indicates the liquid-liquid phase boundary. **(b)** Fluorescence recovery after photobleaching (FRAP) plots of FUS (10 μM) condensates at various concentration of PEG8000. **(c)** Confocal fluorescence microscopy images corresponding to the data in Fig. 1b are shown. Scale bar = 4 μm. **(d)** Analysis of the FRAP results from Fig. 1b reveals scaling of apparent diffusion coefficients *(left axis; blue line)* and the fraction of mobile phase *(right axis; orange line)* with increasing concentration of PEG8000.

The observed activity of PEG on lowering the RNP liquid-liquid coexistence boundary can be explained by a crowder-mediated depletion force that effectively increases the net inter-RNP attractive potential with increasing concentration of the crowder[52] (also *see* SI note 1). If that is the case, we expect that the fluidity of RNP droplets will decrease with increasing macromolecular crowding due to enhancement of the inter-molecular network strength within the condensed phase[40]. Similar phenomenon is known to drive colloidal gel formation in polymer-colloid mixtures[53]. In case of the RNP condensates, we expect a considerable decrease in the biomolecular diffusion rate with increasing PEG. Therefore, we next investigated the physical properties of FUS condensates as a function of PEG8000 concentration at a fixed protein concentration (10 µM; Fig. **1b-d**). We used two complementary methods to probe FUS^FL^ condensate material states: (a) fluorescence recovery after photobleaching (FRAP), and (b) controlled fusion of suspended droplets using a dual-trap optical tweezer. Using FRAP, we measured the half-time of fluorescence recovery (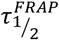) of the fluorescently-tagged RNP (Alexa488-FUS) after photobleaching a circular region at the center of the droplet (Fig. **1b&c**; Fig. **S1**). Analysis of the 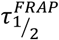 and effective bleach radius (*r*_*e*_) allowed us to estimate apparent diffusion coefficient (*D*_*app*_) of fluorescently tagged RNP within the protein-dense phase (see *SI methods* and Fig. S1). Fraction of mobile phase was also estimated from FRAP data and was used as a relative measure of the viscoelastic properties of FUS droplets[8]. With increasing PEG8000, our FRAP data revealed two distinct trends (Fig. **1b-d**): (i) increase in the fluorescence recovery half-time, and hence, a concomitant decrease in molecular diffusivity, and (ii) decrease in the fraction of mobile phase. Our observed molecular diffusion rate decreased nearly 4-fold upon increasing the PEG8000 concentration from 0 to 150 mg/ml. Concurrently, total fluorescence recovery after bleaching decreased from 100% to ~ 40% (total observation time: 300 s) for FUS^FL^ droplets. These data suggest that macromolecular crowding mediates a progressive transition of FUS droplets from a viscous fluid state to a viscoelastic state resulting in substantially arrested protein diffusion.

To gain further insight into the altered physical properties of FUS^FL^ droplets in presence of macromolecular crowders, we performed quantitative droplet fusion experiments using a dual-trap optical tweezer. In these experiments, we used one laser beam to hold one RNP droplet at a fixed position while another RNP droplet, trapped by a second laser, was moved towards the first trapped droplet at a constant velocity[46] (Fig. **2a**; Fig. **S2**). As these droplets were brought into proximity, liquid FUS^FL^ droplets coalesced rapidly in the absence of crowders with a timescale of ~ 200 ms/µm (Fig. **2**, *top panel*; Supplementary movie 1). In presence of 150 mg/ml PEG8000, the fusion events of FUS^FL^ droplets were almost arrested (incomplete fusion in Fig. **2**, *bottom panel*; Supplementary movie **3**). Instead, we observed that FUS^FL^ droplets cluster at the optical trap under this condition (Fig. **S3**). This is consistent with our FRAP results (Fig. **1b-d**). Together, our observations from FRAP and optical trap experiments lead us to conclude that macromolecular crowding by PEG substantially alters the physical and material properties of FUS^FL^ droplets.

**Figure-2:**
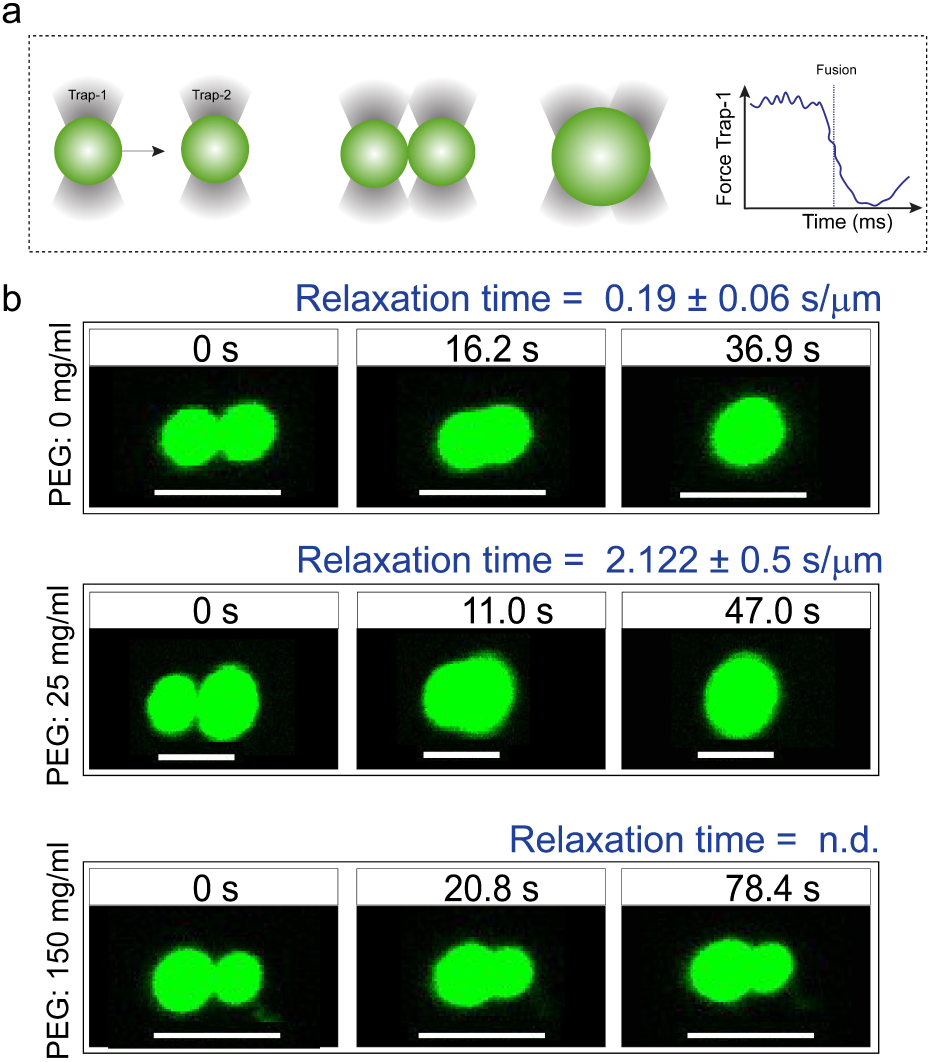
Coalescence dynamics of suspended condensates provide insights into FUS^FL^ droplet material states. **a.** Experimental scheme using a dual-trap optical tweezer. **b.** Controlled fusion of suspended FUS^FL^ droplets by the optical tweezer in the presence of 0 (top), 25 (middle), and 150 (bottom) mg/ml of PEG8000. The normalized relaxation times are indicated in each case. For 150 mg/ml PEG samples, droplets did not completely fuse. Scale bar = 5 μm. n.d.: not determined. Also see *Supplementary movies 1-3.*

### The effect of macromolecular crowding on RNP condensation is independent of LCD sequence

The phase separation of FUS family of RNPs is assumed to be driven by enthalpy, i.e., attractive protein-protein interactions, and therefore, is sensitive to LCD sequence features that encode essential intermolecular interactions[7]. As such, several recent studies were focused on elucidating the sequence determinants of LLPS in RNPs, which revealed two major classes of LCDs[45, 46]. These are (i) prion-like LCD (PrD), characterized by an overabundance in polar (S/G/Q) and aromatic (Y/F) residues but largely devoid of charged amino acids[1, 2, 7, 8, 40, 54-56], and (ii) arginine-rich polycationic LCD (R-rich LCD)[57], such as the disordered RGG-box sequences present in RNA-binding domains of many RNPs[58]. In the ribonucleoprotein FUS, both of these LCD types (FUS^PrD^ and FUS^RGG^) are present as individual domains in the N- and C-termini of the protein, respectively (Fig. **3a**; Supplementary Table-1). To gain mechanistic understanding of the impact of crowding on the physical properties of RNP droplets formed by the distinct LCDs, we next studied the phase behavior and condensate property of FUS^PrD^ and FUS^RGG^, independently. FUS^PrD^ is intrinsically disordered, as predicted bioinformatically[59] (Fig **3a**) and verified experimentally[48]. Interestingly, FUS^PrD^ is also known to form amyloids *in vitro* that contain dynamic β-sheet rich structures[60, 61], which is assumed to facilitate the “droplet aging” process[41]. While FUS^PrD^ has been implicated as the major driver of FUS LLPS[62], the role of FUS^RGG^ in RNP condensation is only beginning to be explored[51]. Similar disordered RGG-box regions with low complexity R/G rich sequence motifs have previously been shown to form homotypic condensates[63]. Here, to study the effect of crowding, we first constructed phase diagrams of PEG-LCD mixtures for these two LCDs separately. Both LCD types underwent concentration dependent reversible phase separation upon lowering the solution temperature (Fig. **S4**), suggesting that there exists an upper critical solution temperature (UCST) for these two disordered domains above which a homogeneous phase is energetically favored. The observed UCST phase behavior for two LCDs also indicates that their phase separation is driven predominantly by favorable free energy change during self-association via attractive protein-protein interactions[7]. We observed that PEG8000 facilitates phase separation of both FUS LCDs. Isothermal phase diagram (at 22±1 ^°^C) analyses for FUS^PrD^ and FUS^RGG^ as a function of PEG8000 are presented in Fig. **3**. We observed that PEG decreases the protein condensation critical concentration by ~ 5 fold for FUS^PrD^ in response to an increase in PEG concentration from 0 to100 mg/ml. A similar trend is also observed for FUS^RGG^, where the concentration required for the protein to undergo LLPS is decreased by ~ 7 fold in presence of 100 mg/ml PEG. These results are consistent with the hypothesis that the crowder mediated depletion force acts synergistically to enhance the homotypic LCD-LCD interactions (see SI note 1).

**Figure-3:**
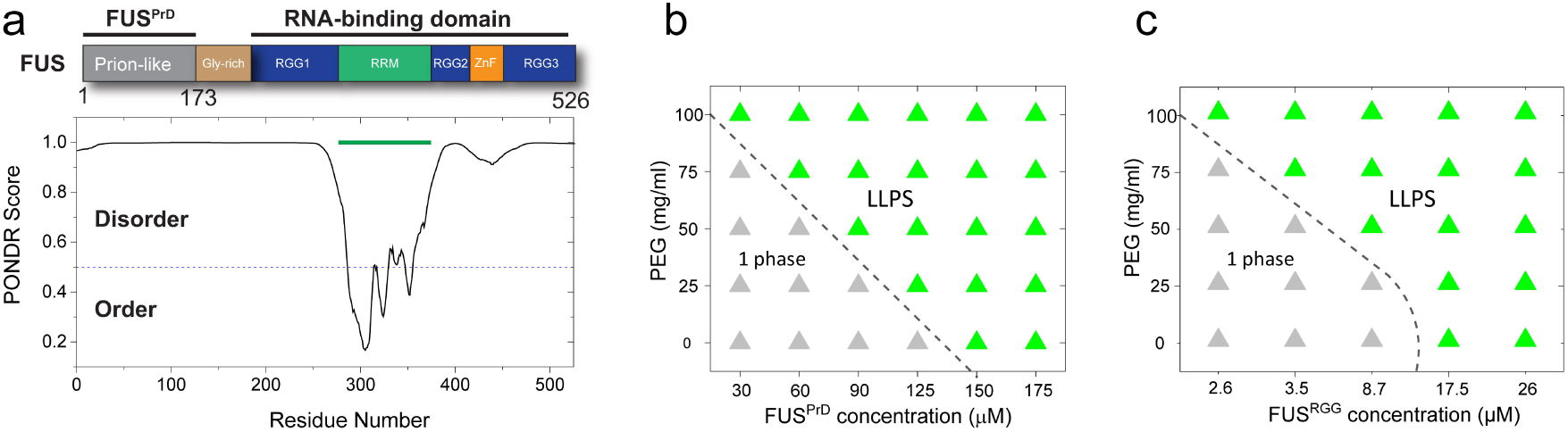
FUS harbors distinct low-complexity disordered domains (LCDs) that individually undergo crowding-mediated LLPS. **a.** Domain architecture of FUS; two distinct disordered domains are highlighted. The N-terminal FUS LCD has prion-like sequence features (FUS^PrD^) and C-terminal LCD is enriched in RGG-repeats (FUS^RGG^). The disorder prediction scores (using VSL2 algorithm^59^) are also shown. **b&c.** Effect of PEG on the LLPS of prion-like LCD and R-rich LCD. Shown here are the phase diagram of FUS^PrD^ and FUS^RGG^ with PEG8000, respectively. The dotted lines indicate the liquid-liquid coexistence boundaries.

During the visualization of FUS^PrD^ and FUS^RGG^ condensates using a confocal fluorescence microscope, we noted that the well-dispersed LCD droplets undergo clustering at higher PEG concentrations (Fig. **S5**). This observation indicates a crowding-dependent change of condensate physical properties. Therefore, we next investigated LCD droplet material properties in the presence of macromolecular crowders. Using Alexa488-labeled LCDs, we quantitatively analyzed fluorescence recovery after photobleaching in a well-defined region within FUS^PrD^/ FUS^RGG^ droplets (Fig. **4**). The recovery of fluorescence is modelled based on the diffusion of LCDs within the respective condensed phases (Fig. **S1**). For FUS^PrD^ droplets, FRAP analysis reveal that the diffusion rate decreases significantly with increasing the PEG8000 from 0-175 mg/ml. Only ~15% fluorescence recovery was observed at ≥ 150 mg/ml PEG8000 in a timescale of 300 s after bleaching (Fig. **4a-c**). The FUS^RGG^ droplets showed a similar transition from a liquid state to a viscoelastic state by PEG8000 (Fig. **4d-f**), although the changes were observed to be more gradual as compared to FUS^PrD^. These data suggest that both FUS^PrD^ and FUS^RGG^ droplets undergo progressive hardening with increasing PEG, despite their diverse sequence features and charge patterning.

**Figure-4:**
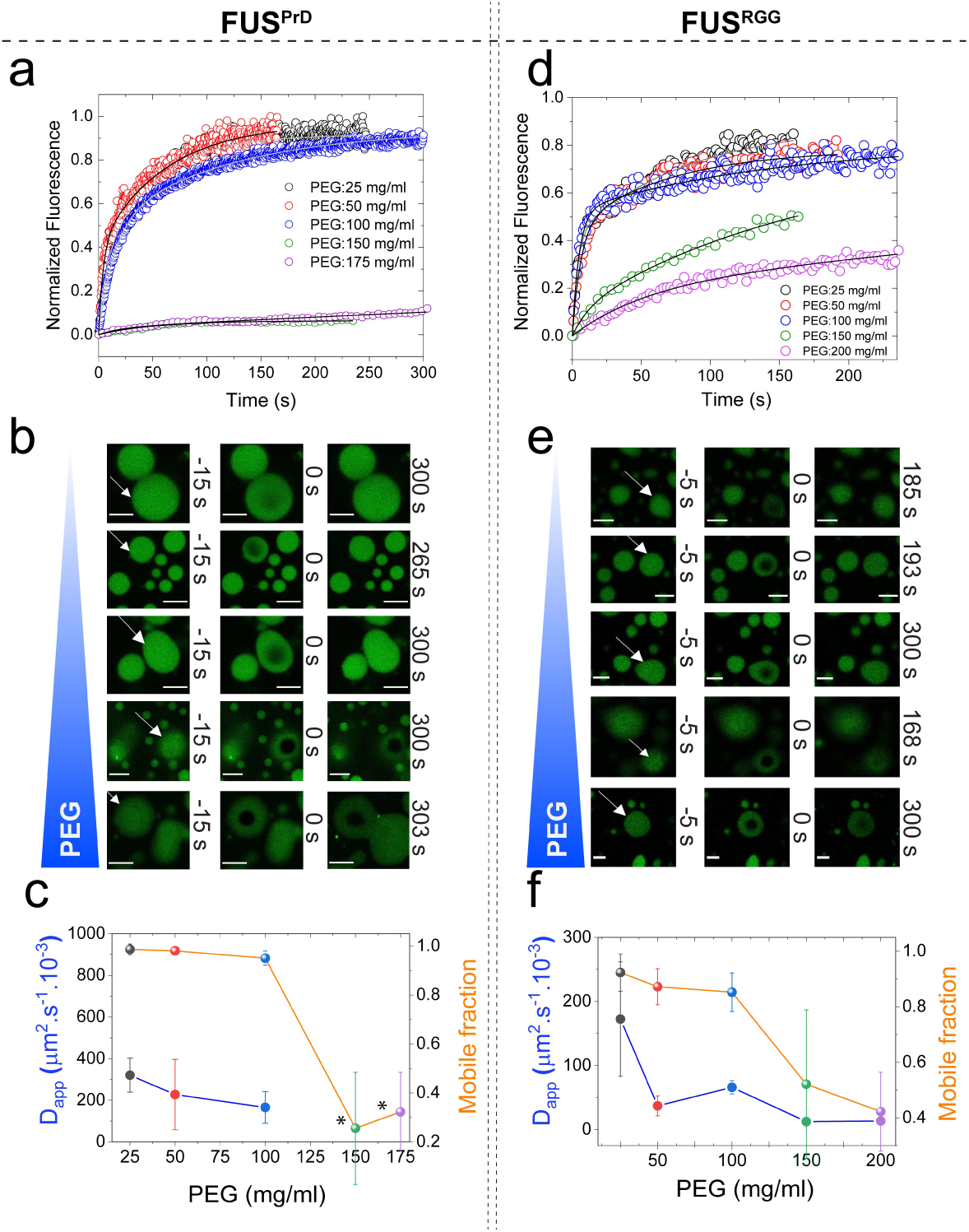
Macromolecular crowding tunes viscoelastic properties of both prion-like and R-rich LCDs. a,b,d,e. Representative FRAP plots and images of FUS^PrD^/FUS^RGG^ droplets at variable concentration of PEG8000, respectively. Scale bar = 8 μm in **b** and = 4 μm in **e**. **c&f.** Analyses of the FRAP data estimating apparent diffusion coefficients of the LCD within the condensed phase *(left axis; blue)* and the mobile phase fraction *(right axis; orange)* in respective cases. Due to a very low fraction of recovery, D_app_ estimation from the FRAP data for FUS^PrD^ droplets at 150 and 175 mg/ml PEG was omitted (indicated by the asterisks in *c*).

### Crowding impact on the material properties of FUS condensates is observed for a broad range of crowders

Macromolecular crowding in a cell arises from biopolymers with a plethora of sizes and shapes. Therefore, to evaluate the excluded volume effect on RNP LLPS, it is necessary to use polymer crowders with variable chain length[25, 64]. Therefore, we next considered the impact of crowders with different molecular weights on FUS^FL^ condensation. To this end, we employed PEG polymers with molecular weights ranging from 300 gm/mol to 35,000 gm/mol. FRAP experiments reveal that FUS^FL^ droplets are viscoelastic in the presence of all the crowders tested (crowder concentration = 150 mg/ml), with a significant degree of arrest in the dynamic exchange of the fluorescently tagged RNP (Fig. **5a**; Fig. **S6a**). Dextran, another widely utilized molecular crowder with a different chemical identity, also showed similar effects on FUS^FL^ condensates (Fig. **5b**; Fig. **S6b**). These data collectively suggest that our observed viscoelastic tuning of FUS condensates is generally applicable to a broad range of polymer crowders and, therefore, represents a common effect of deletion interaction as induced by macromolecular crowding.

**Figure-5:**
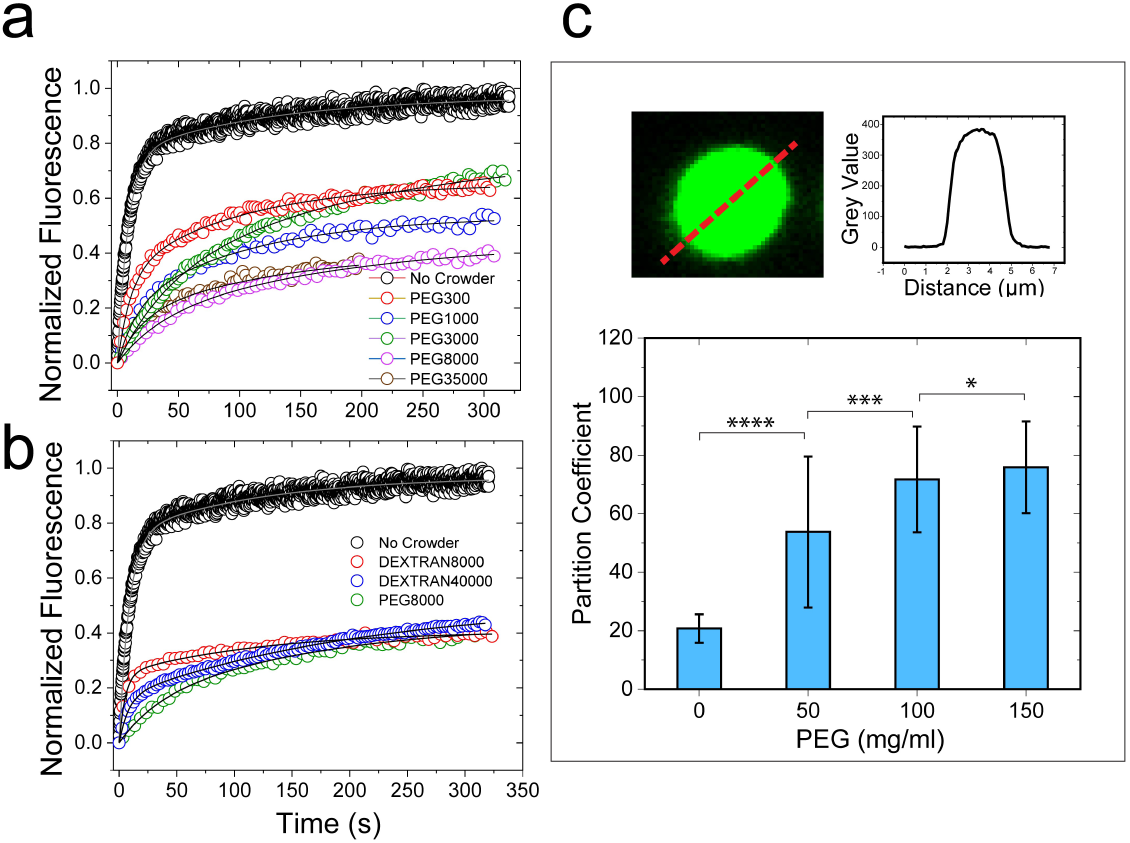
PEG and dextran produce similar effects on FUS^FL^ droplet physical properties. **a&b.** Representative FRAP traces at 150 mg/ml PEG/dextran with a wide range of molecular weight, as indicated. The corresponding FRAP images and diffusion analyses are shown in Fig. S6. **c.** Increased RNP partitioning within the FUS^FL^ droplets with increasing concentration of PEG8000. * p-value: 0.1-0.01, ** p-value: 0.01-0.001, *** p-value: 0.001-0.0001, **** p-value < 0.0001.

## Discussion

Intracellular RNP granules are phase separated bodies that display characteristic dynamics of complex fluids[2, 65]. The physical properties of RNP condensates are important modulators of their functions[3]. Aberrant alterations of the droplet material states, such as age onset loss of granule fluidity and formation of solid-like mesoscale assemblies, are implicated in various neurological disorders including ALS and FTD[41, 45, 49]. Over the past two years, considerable efforts have been dedicated in characterizing RNP sequence-encoded molecular interactions that control the material properties of the RNP condensates[51]. In this study, we consider the role of molecular crowding, a ubiquitous thermodynamic force in the cellular environment, on the RNP condensate dynamics. We experimentally show that crowding, as mimicked by biocompatible “inert” polymers, not only lowers the LLPS boundary, but also substantially alters the exchange dynamics of the RNP within the condensed phase. We observe that the effect of crowding on LCD-driven LLPS is largely independent of respective disordered domain sequences. To provide a mechanistic picture of the observed effect of crowding on the RNP condensation, we consider a thermodynamic model that describes the perturbation of protein-protein interactions by a crowder in light of the well-established excluded volume effect[30, 33]. According to this model, introduction of a polymer crowder in the RNP-buffer solution leads to an isotropic inter-protein attraction by the depletion force (*see* SI note 1), which acts in tandem with the intrinsic LCD-LCD homotypic attraction. The physical origin of the depletion attraction is the exclusion of the center of mass of a crowder molecule from a region surrounding an RNP molecule, which is typically called the depletion layer (Fig. **6**). The depletion layer thickness is directly proportional to the size of the polymer crowder[34]. A salient feature of this simple model is — to produce excess free volume available to the polymer crowders, increasing crowder concentration in the solution increases the overlap of the RNP depletion layers (Fig. **6**). This implies that the net attractive LCD-LCD attraction is enhanced by macromolecular crowding, which lowers the critical concentration of RNP LLPS and results in hardening of RNP condensates (SI note 1). One key prediction of this model is that RNP partitioning in the dense phase should increase with increasing crowding (Fig. **S7**). This was experimentally verified for FUS^FL^ condensates using confocal fluorescence image analysis, which indicates that the RNP partition increases by ~ 4 fold in response to an increase in PEG8000 concentration from 0 mg/ml to 150 mg/ml (Fig. **5c**)

**Figure-6:**
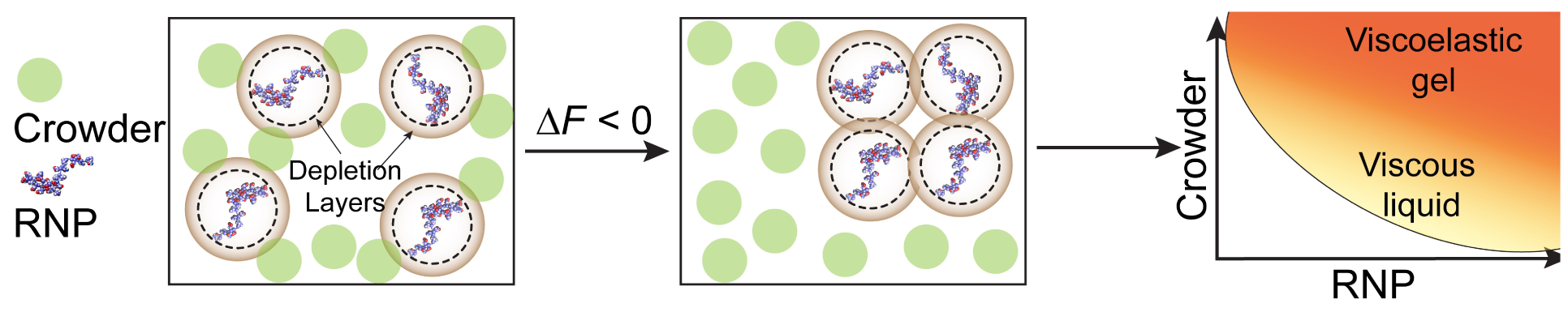
Schematic representation of depletion-attraction: Proposed model showing crowder-mediated overlap of depletion layers as a driving force underlying RNP droplet formation and maturation into a gel-like phase.

In summary, we demonstrated that depletion interaction, as induced by macromolecular crowding, continuously tunes the physical properties of RNP condensates ranging from purely viscous fluids to viscoelastic gel-like states. Alteration in the material properties of phase separated membraneless compartments inside cells has been previously observed both in normal physiology and pathology[41, 66, 67]. Our results suggest that the hardening of RNP condensate is considerably influenced by the entropic forces in crowded cellular environment. Several recent reports focused on identifying key RNP sequence features and role of RNA/protein partner binding that contribute to the gelation of RNP droplets[10]. Based on the data presented here, we postulate that generalized thermodynamic forces that can tune effective RNP homotypic interactions are likely to influence the rate at which physiological condensates mature into a viscoelastic gel.

## Acknowledgements

The authors gratefully acknowledge UB north campus confocal imaging facility (supported by National Science Foundation MRI Grant: DBI 0923133) and its director, Mr. Alan Siegel for helpful assistance.

## Author contributions

P.R.B. designed the study. T.K., I.A., and P.R.B. designed the experimental strategies with occasional input from M.M.M. W.W. expressed and purified recombinant proteins from bacteria. T.K., W.W., and P.R.B. performed the phase diagram analysis. T.K. and I.A. collected confocal microscopy experiments, partitioning, FRAP measurements, and trap-induced droplet fusion with help from P.R.B. T.K., I.A., and J.N. analyzed all the data. P.R.B., T.K., and I.A. wrote the manuscript.

