## Supplementary material for "Molecular Crowding Tunes Material States of Ribonucleoprotein Condensates": Materials and Methods, Supplementary table-1; Supplementary Figures-S1-S7; Supplementary Note-1; Supplementary Movie Legends

*Protein Samples:* Codon optimized wild-type full-length FUS ($FUS^{FL}$), prion-like domain of FUS ($FUS^{PrD}$; AA: 1-173), and the RNA-binding domain of FUS without the zinc-finger domain ($FUS^{RGG}$:211-526Δ422-453) were gene synthesized by GenScript USA Inc. (NJ, USA) and cloned into (cloning site: SspI-BamHI) pET His6 MBP N10 TEV LIC cloning vector (2C-T). The plasmid vector was a gift from Scott Gradia (Addgene plasmid # 29706). TEV-cleaved proteins contained three exogeneous amino acids (SNI) at its N-termini. *E. Coli* cells (BL21(DE3)) were transformed with the plasmids containing $FUS^{FL}$ and its variants in respective cases. Transformed cells were induced with IPTG (0.5 mM final concentration) at OD = 0.6 - 0.8 and further grown for an additional 3 - 5 hrs at 30 °C. Protein extraction was performed using a french press in lysis buffer (50 mM Tris-HCl, 10 mM imidazole, 1 M KCl, pH 8.0) containing protease inhibitor cocktail (Roche). Cell debris were removed by centrifugation. His-tag proteins were purified from the crude cell lysate using Ni-NTA agarose matrix (Qiagen Inc, Valencia, CA) by gravity-flow chromatography following the manufacturer's protocol with the following modifications: the wash buffer included 1.5 M KCl to disrupt nucleic acid binding to the recombinant protein[1,2], which was eluted with elution buffer containing 250 mM imidazole and 150 mM NaCl. The purity of the eluted protein samples was checked using $A_{280}/A_{260}$ measurements (to rule out presence of nucleic acids), and by polyacrylamide gel electrophoresis (PAGE) and Coomassie blue staining. The eluates (individual or pooled) were dialyzed against 25 mM Tris-HCl, pH 7.5 buffer containing 10% glycerol. The concentration of the protein samples were determined by absorbance at 280 nm using the following extinction coefficients: 103,600 $M^{-1}$ $cm^{-1}$ for $FUS^{PrD}$-MBP, 86,750 $M^{-1}$ $cm^{-1}$ for $FUS^{RGG}$-MBP, and 138,000 $M^{-1}$ $cm^{-1}$ for $FUS^{FL}$-MBP (https://web.expasy.org/protparam). The protein samples were flash frozen in small aliquotes and stored in -80 °C.

*Fluorescence labeling:* The S86C variant of $FUS^{PrD}$ and A313C variant of the $FUS^{RGG}$ were expressed and purified using an identical protocol as described above for the WT protein, except one modification: all buffers contained 2 mM DTT. The protein samples were fluorescently labeled with Alexa488 dye (C5-maleimide derivative, Molecular Probes) using Cys-maleimide chemistry as described in our earlier work[3]. The labeling efficiency for all samples were observed to be ≥ 90% (UV-Vis absorption measurements), and no additional attempt was made to purifiy them further, given that only labeled protein is observed in the fluorescence experiments.

*Sample preparation for phase separation measurements:* All of the protein samples were buffer exchanged into the phase separation buffer (25 mM Tris-HCl, pH 7.5) containing 150 mM NaCl unless otherwise noted. Prior to performing phase separation measurements, the $His_6$-MBP-N10 tag was removed by the action of TEV protease (1:25 ratio) (GenScript USA Inc.) for 1 hr at 30 °C. The completion of the cleavage reaction was judged by polyacrylamide gel electrophoresis (PAGE) and Coomassie blue staining.

*Phase diagram analysis:* Phase diagrams were constructed by turbidity measurements at 350 nm using a NanoDrop oneC UV-Vis spectrophotometer at room temperature (22±1 °C). Desired amounts of PEG solutions were added to the protein solutions from a 35% (w/v) stock in nuclease-free water with appropriate salt concentrations. Each sample was incubated ~ 120 seconds prior to turbidity measurements using a 1 mm optical path length. Simultaneously, visualization of protein droplets (or lack thereof) was performed using a Primo-vert inverted iLED microscope (Zeiss), equipped with a Zeiss Axiocam 503 monochrome camera. A global analysis of the turbidity and microscopy data was performed to construct phase diagrams in respective cases based on a simple binary criterion that identifies if droplets were present at a given protein/PEG concentration.

*Confocal fluorescence microscopy:* The fluorescence and DIC imaging were performed using a Zeiss LSM 710 laser scanning confocal microscope, equipped with a 63x oil immersion objective (Plan-Apochromat 63x/1.4 oil DIC M27) and a Zeiss Primovert inverted microscope. Samples were prepared and imaged using tween-coated (20% v/v) Nunc Lab-Tek Chambered Coverglass (ThermoFisher Scientific Inc.) at room temperature (22±1 ºC) unless otherwise noted, with ~ 1% labeled protein samples within the mixture of unlabeled proteins. All the samples were allowed to equilibrate in the chambered coverglass for ~30-45 mins before imaging. For Alexa488-labeled samples, the excitation and emission wavelengths were 488 nm/503-549 nm. Fluorescence recovery after photobleaching (FRAP) experiments were performed using the same confocal set up. The images and data were analyzed using Fiji software[4] and the FRAP curves were plotted and analyzed using origin software (OriginPro 2018).

*Fluorescence recovery after photobleaching (FRAP):* FRAP experiments were performed using Zeiss LSM 710 laser scanning confocal microscope as described above. A circular region of interest (ROI) was bleached with 2-5 iterations of scanning using 100 % laser power for a total time of 2-18 s. Fluorescence intensity changes with time were recorded for three different ROIs (bleached droplet, reference droplet, and background) for approximately 300 s or until the bleached ROI recovered and reached an equilibrium state. Data analyses were performed using Fiji software and MATLAB.

The fluorescence intensities from bleached ROI were corrected for photofading by multiplying with a correction factor obtained from reference ROI as follows:

$$C_f = \frac{R_i}{R(t)}$$

$$I_{corrected} = C_f \times I_{bleached}(t)$$

$R_i$: Initial intensity of reference droplet
$R(t)$: Intensities of reference ROI at time *t*
$C_f$: Correction factor
$I_{bleached}(t)$: Intensity of bleached ROI at time *t*
$I_{corrected}(t)$: Intensity of bleached ROI (corrected for photofading) at time *t*

The corrected itensities were shifted to set the immediate post-bleach point to zero.

$$I_{shifted}(t) = I_{corrected} - \text{min. value of } I(t)$$

which were then normalized.

$$I_{Normalized}(t) = \frac{I_{shift}\ (t)}{max.\ value\ of\ I_{shifted}(t)}$$

This normalized intensity post-bleach was plotted *vs.* time and fitted with a single exponential $y = A\,(1 - \exp(-t/\tau)$ using MATLAB. Half time of recovery ($\tau_{1/2}$) was obtained from the fitting parameter. To improve the goodness of the fit, two-exponential fit $y = A\,(1 - \exp(-t/\,\tau_A) + B\,(1 - \exp(-t/\,\tau_B)$ was also used.[5] Half time of recovery ($\tau_{1/2}$) was obtained graphically for the latter (Fig. S1c,d).

To account for diffusion during bleaching, instead of simply using user defined bleach radius ($r_n$) for diffusion coefficient calculations, an effective radius ($r_e$) from the immediate post bleach frame was calculated by taking a profile across bleached ROI in Fiji. The normalized fluorescence intensities were plotted with distance and fitted with an exponential of a Gaussian laser profile using MATLAB, as previously described.[6,7]

$$f(x) = \exp(-Kexp\left(\frac{-2(x-b)^2}{{r_e}^2}\right) \tag{1}$$

$r_e$ obtained from the fitting is the effective radius which corresponds to half width at 86 % of bleach depth K (Fig. S1a,b).

The apparent diffusion coefficient[6] was calculated using the following equation:

$$D = \frac{{r_e}^2 + {r_n}^2}{8\tau_{1/2}} \tag{2}$$

The mobile fraction[6,7] was calculated using the following formula:

$$M_f = \frac{I_\infty - I_0}{I_i - I_0} \tag{3}$$

$I_\infty$: Fluorescence intensity after recovery
$I_0$: Fluorescence intensity immediately after bleach
$I_i$: Fluorescence intensity before the bleach

*Coalescence of suspended droplets by dual-trap optical tweezer:* Controlled fusion assays were conducted to investigate changes in the material properties of the $FUS^{FL}$ condensates as a function of crowder concentration. The samples were injected into a 25 mm x 75 mm x 0.1 mm single chamber custom-made flow cells. Samples at 10 µM protein concentration were prepared with different PEG8000 concentrations and equlibrated for ~ 30 minutes at room temperature. The induced fusion was done using a dual-trap optical tweezer system coupled with laser scanning confocal fluorescence microscope (LUMICKS™ C-trap). In a typical fusion experiment, two droplets were trapped with a 1064 nm laser with minimum power to reduce heating effect. The trapping of droplets was acheived due to a difference in the refractive index between the condensate and the dilute phase. After trapping, one droplet is brought into contact with the other droplet at a constant velocity of 40 nm/s. The trap remains traveling at that velocity until the fusion is completed and the final droplet relaxes to a spherical shape. The force on the moving trap was recorded at 78.4 kHz sampling frequency and analyzed using a fusion relaxation model[8]. The following equation was used to fit the force-time curve:

$$F = ae^{(-t/\tau)} + bt + c \tag{4}$$

Where the parameter $\tau$ is the fusion relaxation time. We scaled the fusion time by the average of the radii of the two droplets for every event. The linear term in the model is added to account for the constant trap velocity. First, we did a control experiment on a slow fusion sample and the relaxation times were obtained both from force curves as well as from aspect ratio analysis using fluorescence images[9]. The results were in good quantitative agreement (data not shown). For $FUS^{FL}$ samples, at least 15 droplet fusion events were collected for each PEG concentration and the scaled relaxation times were averaged. An example of a typical normalized force curve with the fitted model is shown in Fig. S2.

*Partition analysis:* Phase separated samples containing appropriate amount of fluorescently tagged protein were placed in a single-chambered custom-made flow cell (see *coalescence of suspended droplets* section). Droplets were imaged at the surface using laser scanning confocal fluorescence microscope using 60x water-immersion objective (LUMICKS™, C-trap). Images were analyzed using Fiji software. To calculate the partition coefficient, the mean intensity of the entire droplet was divided by mean background intensity for several droplets per sample for statistical accuracy using Excel. Statistical analysis were carried out using MATLAB.

**Supplementary Table-1**

| Construct Name | Amino acid sequence |
|---|---|
| $FUS^{PrD}$ | MASNDYTQQATQSYGAYPTQPGQGYSQQSSQPYGQQSYSGYSQSTDTSGYG QSSYSSYGQSQNSYGTQSTPQGYGSTGGYGSSQSSQSSYGQQSSYPGYGQQP APSSTSGSYGSSSQSSSYGQPQSGSYSQQPSYGGQQQSYGQQQSYNPPQGYG QQNQYNSSSGGGGGG GGG |
| $FUS^{RGG}$ | QDRGGRGRGGSGGGGGGGGGGYNRSSGGYEPRGRGGGRGGRGGMGGSDR GGFNKFGGPRDQGSRHDSEQDNSDNNTIFVQGLGENVTIESVADYFKQIGIIK TNKKTGQPMINLYTDRETGKLKGEATVSFDDPPSAKAAIDWFDGKEFSGNPI KVSFATRRADFNRGGGNGRGGRGRGGPMGRGGYGGGGSGGGGRGGFPSGG GGGGGQQGPGGGPGGSHMGGNYGDDRRGGRGGYDRGGYRGRGGDRGGFR GGRGGGDRGGFGPGKMDSRGEHRQDRRERPY |

### Supplementary Figures

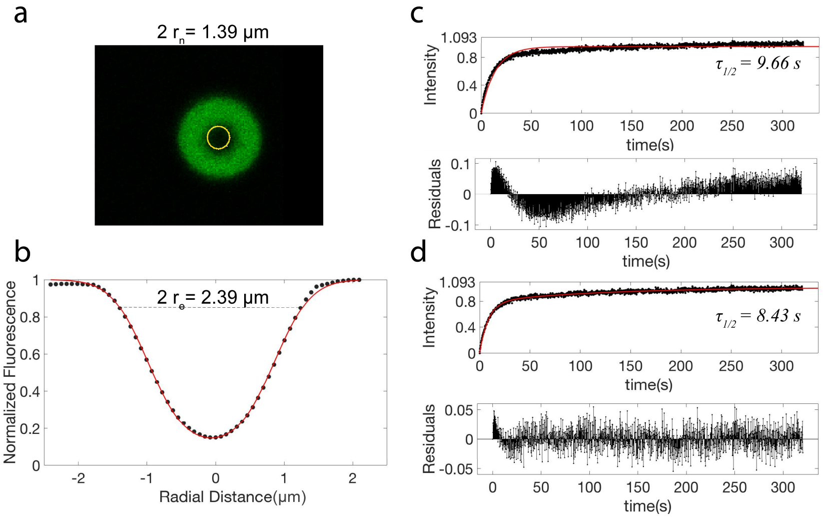


**Figure S1: Determination of biomolecular diffusion by fluorescence recovery after photobleaching (FRAP). a**) Immediate post bleach frame with user defined ROI (yellow circle; $r_n$ = nominal radius). **b**) Normalized intensity profile across the bleached ROI to estimate the effective radius of bleaching ($r_e$). Black points are the data, red line is a fit to an exponential of a Gaussian laser profile using equation-1. **c**) Normalized FRAP curve (black points) fitted with a single exponential funcion (red line). **d**) Normalized FRAP curve (black points) fitted with a double exponential function (red line). The residuals are shown in respective cases, justifying our choice of a double exponential function to determine the $\tau_{1/2}$.

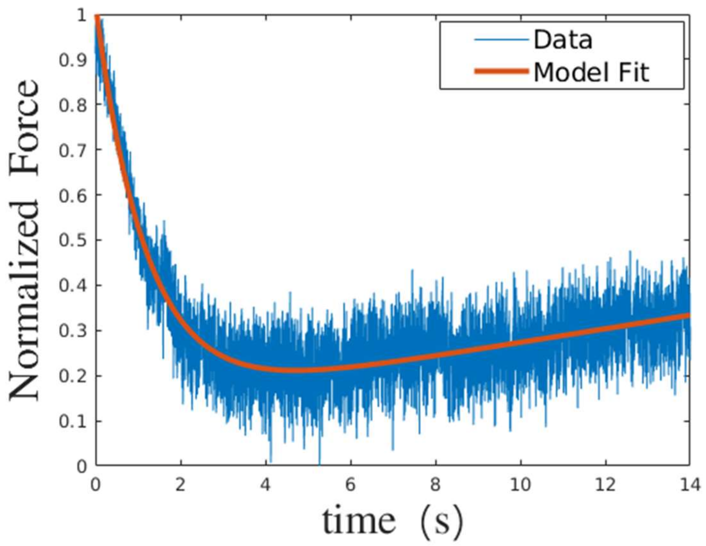


**Figure S2: Representative normalized force relaxation curve during trap-induced coalescence of suspended FUS$^{FL}$ droplets**. The data (blue trace) is fitted (red line) using a model described by equation-4.

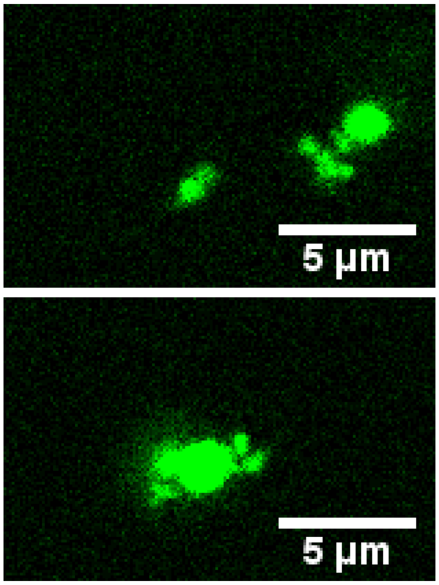


**Figure S3: Aggregation of FUS$^{FL}$ condensates in the optical trap in presence of 150 mg/ml PEG8000**. Two images (top and bottom) correspond to two independent measurements. The green color corresponds to Alexa488-labeled FUS$^{PrD}$. FUS concentration = 10 µM.

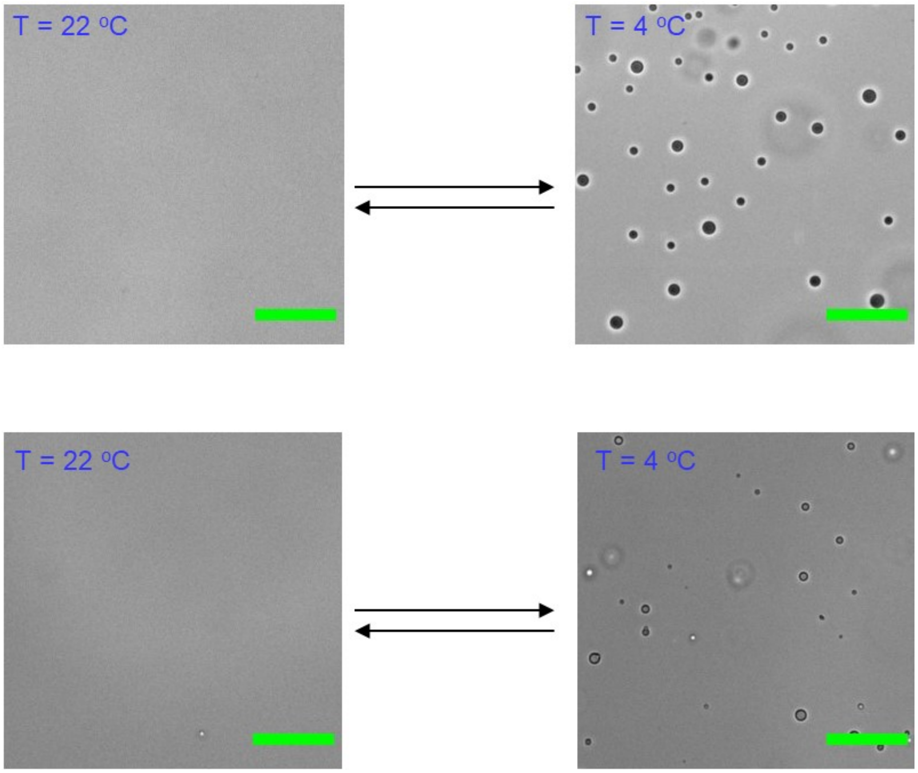


**Figure S4: FUS$^{LCD}$ condensation is reversible and exhibit upper critical solution temperature (UCST).** Effect of temperature on FUS$^{PrD}$ (100 µM; *top two panels*) and FUS$^{RGG}$ (25 µM; *bottom two panels*) LLPS, revealing UCST phase behavior. The brightfield images were collected using protein samples in 25 mM Tris.HCl buffer, pH 7.5, 150 mM NaCl. Scale bar = 25 µm.

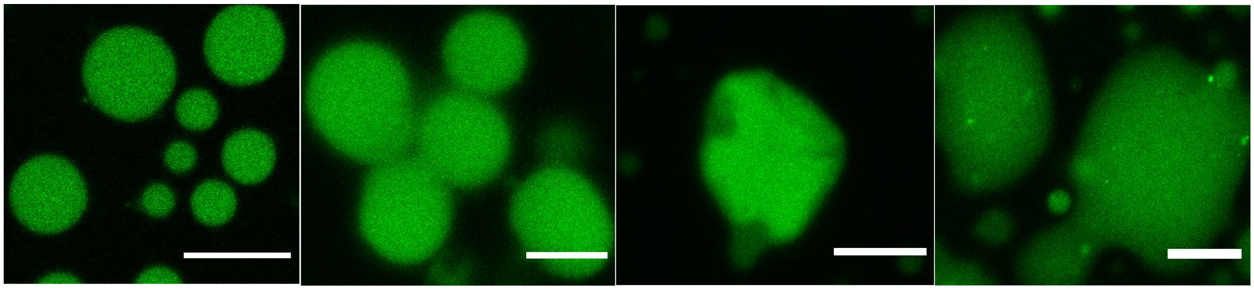


**Figure S5: Clustering and morphological changes of $FUS^{PrD}$ droplets with increasing concentration of PEG8000.** PEG concentrations are indicated in the figure. Protein concentration = 335 µM. Scale bar =10 µm.

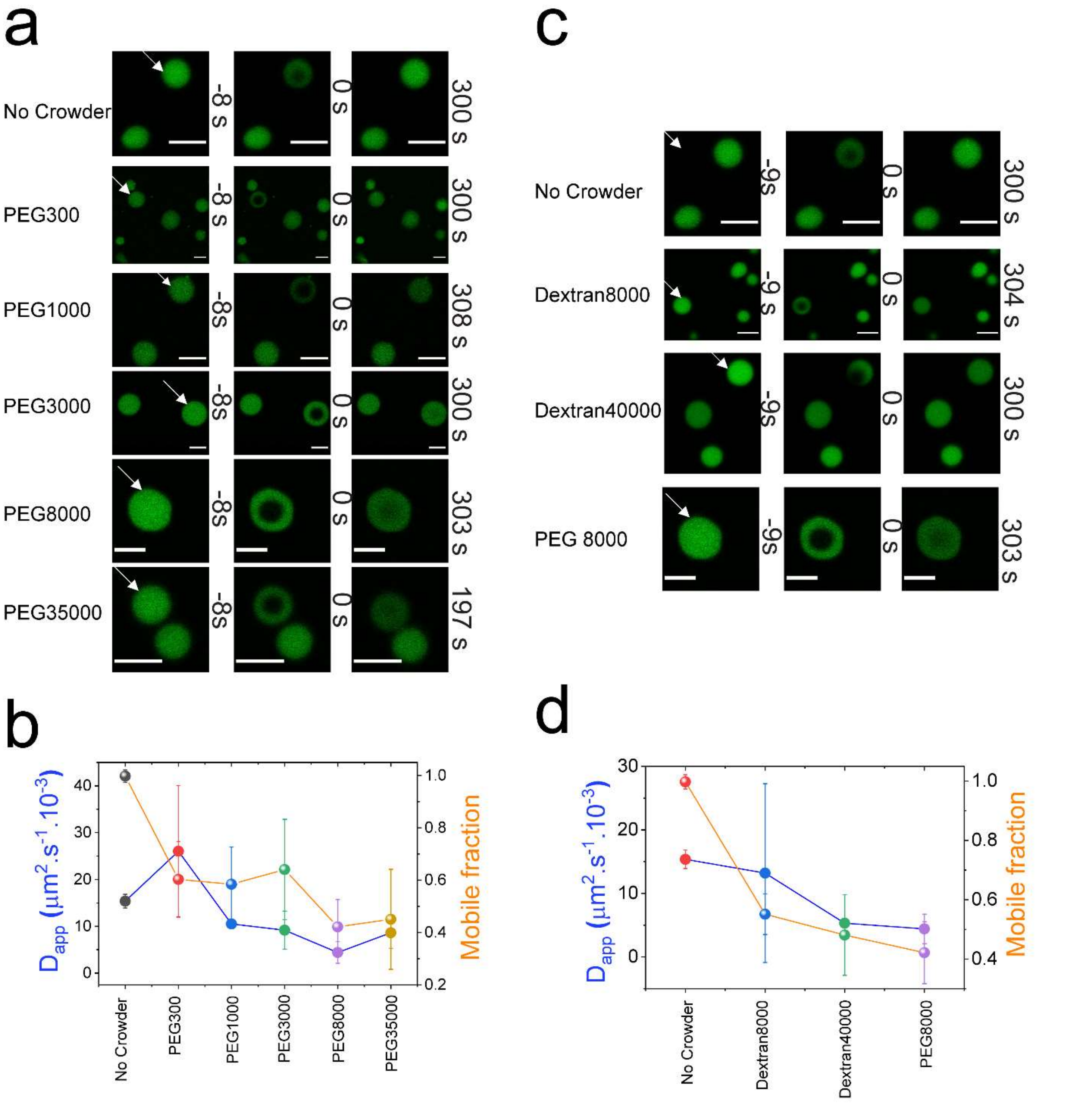


**Figure S6: FRAP analysis of FUS droplets in presence of various crowders**. **a&c.** Representative FRAP images of FUS droplets in presence of PEG and dextran as polymer crowders with different molecular weights of PEG and dextran, respectively. Crowder concentration used in each case: 150 mg/ml. Scale bar =4 μm. **b&d.** Corresponding analysis of the FRAP results to quantify molecular diffusion (blue; *right axis*) and the fraction of mobile phase (orange; *left axis*).

**SI Note-1**

**A thermodynamic model describing the effect of a polymer crowder on the LLPS of FUS**

To describe the effect of a polymer crowder on the LLPS of FUS-buffer system and to analyze the impact of crowder concentration on the effective FUS-FUS interactions, we consider a thermodynamic perturbation theory that was previously developed for gammaD crystallin-buffer-PEG mixture[10,11]. Using the subscripts "pro" to denote the protein and "pol" to denote the polymer crowder, we define the reduced free energy of the system: $f = (F - F_0)/RTV$; F and $F_0$ are Helmholtz free energy of FUS-crowder-buffer system and the buffer system without the protein/polymer crowder, respectively. $F_0$ is also called the standard free energy and is given by: $F_0 = c_{pro}.\mu^0_{pro} + c_{pol}.\mu^0_{pol}$; $\mu^0_{pro}$ and $\mu^0_{pol}$ are standard chemical potentials, R = ideal gas constant.

The first order approximation of the free energy with respect to the polymer crowder concentration can be expressed as[11]:

$$f(c_{pro}, c_{pol}, T) = f(c_{pro}, c_{pol} = 0, T) + c_{pol}.ln\frac{c_{pol}}{e} + c_{pol}(\frac{\partial f}{\partial c_{pol}})_{c_{pro}, c_{pol}=0, T} + \cdots \tag{1}$$

Here the 1st term on the right hand side, $f(c_{pro}, c_{pol} = 0, T)$, is the reduced free energy of the FUS-buffer binary mixture in absence of any crowder. The 2nd term signifies a change in the mixing entropy of the system due to the addition of crowder, whereas the 3rd quantity accounts for the changes in the reduced free energy as a function of crowder concentration. For simplicity, we don't consider the contribution of higher order terms, which is the case for ideal polymer crowders. This form of equation-1 can be used to study the variance of protein-protein attraction by a polymer crowder in light of the well-established excluded volume model[12]. The model is based on the consideration that the center of mass of a crowder molecule is excluded from a region surrounding a protein molecule, which is called depletion layer, due to pure steric forces. In such a case[10,11], a free volume fraction parameter ($\alpha$) can be defined that signifies the fraction of volume available to the polymer crowders, $\alpha = exp(-(\frac{\partial f}{\partial c_{pol}})_{c_{pro}, T})$. This definition helps us to re-write equation-1 in the following form:

$$f(c_{pro}, c_{pol}, T) = f(c_{pro}, c_{pol} = 0, T) + c_{pol}.ln\frac{c_{pol}}{e} - c_{pol}.ln\alpha \tag{2}$$

The reduced protein chemical potential of FUS-crowder-buffer system is given by:

$$\mu_{pro}(c_{pro}, c_{pol}, T) = \left(\frac{\partial f}{\partial c_{pro}}\right)_{c_{pol}, T} = \mu'_{pro}(c_{pro}, T) - \frac{c_{pol}}{\alpha}(\frac{\partial \alpha}{\partial c_{pro}})_T \tag{3}$$

The quantity, $\mu'_{pro}$ is the reduced protein chemical potential of FUS-buffer system in absence of any crowders. Differentiation of equation-3 with respect to $c_{pro}$ allows us to derive an expression of effective thermodynamic protein-protein interactions in presence of a crowder:

$$(\frac{\partial \mu_{pro}}{\partial c_{pro}})_{T, c_{pol}} = (\frac{\partial \mu'_{pro}}{\partial c_{pro}})_T - \frac{c_{pol}}{\alpha}(\frac{\partial^2 \alpha}{\partial {c_{pro}}^2})_T \tag{4}$$

The quantities, $(\frac{\partial \mu_{pro}}{\partial c_{pro}})_{T,c_{pol}}$ and $(\frac{\partial \mu'_{pro}}{\partial c_{pro}})_T$ are related to the FUS-FUS interactions, in presence and absence of a crowder, respectively. They also mathematically define the spinodal surface, which is the boundary between the stable and unstable regions of the FUS-crowder-buffer system (Fig. S7). When, $(\frac{\partial \mu_{pro}}{\partial c_{pro}})_{T,c_{pol}} > 0$, the system is stable as a homogeneous solution, whereas it phase separates into two coexisting liquids if $(\frac{\partial \mu_{pro}}{\partial c_{pro}})_{T,c_{pol}} < 0$. The spinodal boundary is defined as $(\frac{\partial \mu_{pro}}{\partial c_{pro}})_{T,c_{pol}} = 0$. In equation-4; the quantity, $(\frac{\partial^2 \alpha}{\partial {c_{pro}}^2})_T$, describes the change in free volume fraction due to overlap of the adjacent depletion layers. Since free volume increases with increasing overlap of the depletion layers, $(\frac{\partial^2 \alpha}{\partial {c_{pro}}^2})_T > 0$. Equation-4 provides a mathematical basis to rationalize the effect of PEG on FUS-buffer mixture (Fig. S7). Starting with a homogeneous FUS-buffer solution without any crowder, the system is stable and hence $(\frac{\partial \mu'_{pro}}{\partial c_{pro}})_T > 0$. As the concentration of PEG increases, the chemical-potential derivative, $(\frac{\partial \mu_{pro}}{\partial c_{pro}})_T$ decreases and becomes zero (i.e., the spinodal condition) at a sufficiently high concentration of the crowder. Further increase in the crowder concentration leads to a spontaneous separation of the mixture into a dense and a dilute phase due to inherent thermodynamic instability (Fig. S7). The net effect of the crowder here may be physically interpreted as an effective increase in FUS-FUS attraction, manifested by depletion interaction. In case of starting FUS concentration ≥ 2 μM, the system phase separates without any crowder (Fig. 1a) and hence, $(\frac{\partial \mu'_{pro}}{\partial c_{pro}})_T < 0$ to start with. In that case, the relative magnitude of effective FUS-FUS attraction, and hence, the partitioning of FUS molecules in the condensed phase increases with the crowder concentration. This is experimentally verified and shown in Fig. 5c (*main-text*).

In summary, the presented thermodynamic model provides a theoretical rationale for our experimental observations that polymer crowders enhance FUS LLPS and condensed phase material properties. We note that the current model is simplified and representative of ideal polymer crowders in dilute regime. Several factors including crowder-RNP interactions, polymer/RNP chain entanglement, etc. should be carefully considered to develop this model further. One exciting future direction would be to study how the RNP-RNP interactions are scaled with a crowder concentration for individual RNPs with distinct LCD sequences, which we envision will be dependent of the LCD sequence itself.

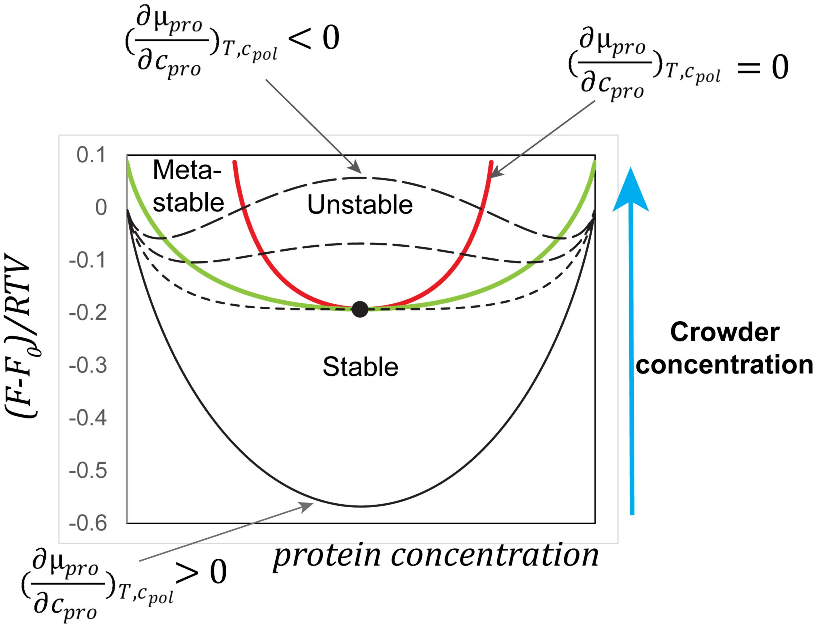


**Figure-S7: Representation of relative free-energy curves as a function of increasing crowder concentration showing transition of a stable homogeneous solution of FUS into a phase separated state.** Shown here are four free energy plots (black-lines). The red line shows the locus of the spinodal line, which separates unstable regions from metastable regions of the mixture. The green line is the coexistence boundary, which specifies the composition of the dense and the dilute phase. With increasing crowding, this model (SI note-1) predicts that the protein partitioning in the dense phase should increase. This is experimentally observed for FUS condensates and shown un Fig. 5c (*main-text*).

**Supplementary Movie Legends**

**Movie-1:** Controlled fusion of suspended $FUS^{FL}$ droplets by a dual-trap optical tweezer in the presence of 0 mg/ml of PEG8000. Scale bar = 5 μm. Speed = 7fps. Duration of each frame = 0.9 s.

**Movie-2:** Controlled fusion of suspended $FUS^{FL}$ droplets by a dual-trap optical tweezer in the presence of 25 mg/ml of PEG8000. Scale bar = 5 μm. Speed = 7fps. Duration of each frame = 0.9 s.

**Movie-3:** Controlled fusion of suspended $FUS^{FL}$ droplets by a dual-trap optical tweezer in the presence of 150 mg/ml of PEG8000. Scale bar = 5 μm. Speed = 7fps. Duration of each frame = 0.9 s.
